# Bayesian Estimation of 3D Chromosomal Structure from Single Cell Hi-C Data

**DOI:** 10.1101/316265

**Authors:** Michael Rosenthal, Darshan Bryner, Fred Huffer, Shane Evans, Anuj Srivastava, Nicola Neretti

## Abstract

The problem of 3D chromosome structure inference from Hi-C datasets is important and challenging. While bulk Hi-C datasets contain contact information derived from millions of cells, and can capture major structural features shared by the majority of cells in the sample, they do not provide information about local variability between cells. Single cell Hi-C can overcome this problem, but contact matrices are generally very sparse, making structural inference more problematic. We have developed a Bayesian multiscale approach, named SIMBA3D, to infer 3D structures of chromosomes from single cell Hi-C while including the bulk Hi-C data and some regularization terms as a prior. We study the landscape of solutions for each single-cell Hi-C dataset as a function of prior strength and demonstrate clustering of solutions using data from the same cell.

## 1 Introduction

The use of whole genome conformation capture techniques (3C) such as Hi-C [10] has revealed that the 3-dimensional (3D) organization of the genome plays a key role in regulating fundamental cellular processes such as transcriptional regulation, cell cycle progression, and cellular differentiation [10, 14, 5]. These studies generate contact maps describing the probability of observing interactions between any two regions of the genome, which can be associated with distance matrices between pairs of genomic loci. Methods developed to infer the 3D structure of chromosomes from these contacts maps typically rely either on optimization-based strategies to minimize the difference between the inferred structure and the distance matrix [21, 8, 28, 32, 17], or on probabilistic modeling to find the most likely structure(s) given the observed contact probabilities [2, 31, 9, 25, 1, 20, 27].

While Hi-C is typically collected on bulk samples containing millions of cells, it is not clear how much the organizational features present in these population datasets reflect the 3D organization of chromosomes in individual cells. For example, it is not guaranteed that all observed long-range contacts appear simultaneously in each cell [26]. Thanks to recent advances in Hi-C technology we can now study long-range interactions at the single cell level [12, 22, 19, 13, 6]. Single cell Hi-C has confirmed many organizational principles described in bulk experiments, but their interpretation is not straightforward. For example, it is not yet clear whether topologically associated domains (TADs) are 3D structural units in individual cells or a population feature that emerges when many cells are aggregated in bulk Hi-C experiments [12, 6], although recent work in *Drosophila* points to the former [24].

The primary difficulty in inferring 3D chromosome structures from single-cell Hi-C data is the sparseness of the contact maps. Currently available methods rely on inference of missing data [18] or on polymer models optimization based on Markov chain Monte Carlo (MCMC) [4] or simulated annealing techniques [12, 22, 13]. However, recovery of potentially missing long range interactions in the contact matrix relies exclusively on the information contained within individual single-cell matrices.

Here we present a solution using Structural Inference via Multiscale Bayesian Approach (SIMBA3D), which utilizes bulk Hi-C to aid in recovering the contribution of interactions potentially missed in single cell Hi-C contact maps. Our strategy is similar in principle to the one used in [26] where bulk Hi-C is decomposed into an ensemble of single cell 3D structures. We build a generalized Bayesian framework which utilizes penalties associated with folding constraints and a prior derived from bulk Hi-C samples to infer 3D chromosomes structure in single cells.

## 2 Method

The primary goal of the inference is to efficiently explore a vast space of potential chromosomal structures and seek optimal solutions using contact matrices and other contextual data. This requires constructing objective functions with desirable properties and developing scalable algorithms to reach interpretable conformations in times that are practical for large scale computations.

As stated above, the problem of estimating chromosomal structure from single cell data is challenging because this data is very sparse and noisy. In order to reach more realistic solutions, we implement a Bayesian approach that supplements the single cell data with the bulk data. This technique helps fill the missing parts with structures corresponding to the population of cells and additionally imposes certain penalties to improve the quality of estimated structures. The penalties are designed in particular to favor uniform placement of points on the estimated curve and to force the curve itself to be smoother.

### Estimation Using Energy Minimization

We define a posterior energy function *E* on the space of potential curves, *X* = ℝ^*n*×3^ – each curve containing *n* points (or nodes) in ℝ^3^ – and use a gradient-based approach to solve for an optimal solution

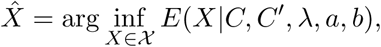
where *C* = (*c_ij_*) and 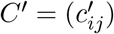 are the single-cell and bulk contact matrices, respectively, and *λ* is a vector of weights. Let 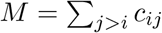 and 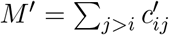, respectively. The energy function *E* has several terms, each contributing to a certain aspect of the estimated curve: 
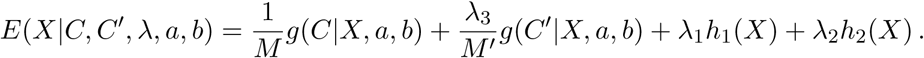

In the first term *g*(*C|X, a, b*) is the negative log-likelihood of the contact matrix *C* given a curve *X*. This term follows a Poisson model (see Varoquaux [28]) with *a*, *b* being pre-determined model parameters. The remaining terms can be viewed as imposing a prior distribution on the curve. The second term involves the negative log-likelihood of the bulk contact matrix *C′* and represents the prior belief that the curve *X* will bear some resemblance to those found in the population. The third and fourth terms represent the prior belief that the curve displays some regularity. The third term penalizes variation in the distances between adjacent points on the estimated curves (the function *h*_1_(*X*) is minimized when the points on *X* are equally spaced), and the fourth term penalizes deviations from straightness (*h*_2_(*X*) is minimized when *X* is a straight line). Since the weights λ_1_ and λ_2_, are typically small, these terms essentially discourage excessive variation in the distances between points and excessive bending of the curve. Together, these terms drive the solution towards a smoother, more interpretable curve that conforms to both single-cell and bulk data. The fact that the gradient of *E* with respect to *X* is available in closed form helps accelerate the search for optimal curves.

Notice that if we let 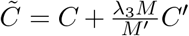 and 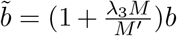, then the Poisson likelihood-based terms *g*(*X|C̃, a, b̃*)*/M+ λ_3_g*(*C′ |X, a, b*)*/M′* merge to become *g*(*X|C̃*, *a*, *b̃*)*/M*. Therefore, the effect of the bulk prior essentially adds a scalar multiple of *C′* to the data *C′* and perturbs the parameter *b*.

### Multiscale Optimization for Improved Inference

The biggest challenge in solving this optimization comes from a non-convex energy function and an extremely high-dimensional search space, which brings about multiple local optima and a tremendous computational cost. While the presence of the bulk and penalty terms helps to mitigate these issues by steering the search towards more realistic solutions, the computational complexity still remains a major hurdle. In order to reduce the computation time to allow for a practical full genome reconstruction, we implement a multiscale optimization technique. Compared to a standard approach that computes the full resolution optimization using a random initialization, the multiscale approach reduces computation time and limits the local solutions obtained. As shown in Fig. 3B, this leads to solutions with smaller energies.

The multiscale optimization technique used in SIMBA3D is as follows. First, from a given full resolution contact matrix, we generate a series of new matrices decreasing in resolution, i.e. decreasing in size, by recursively combining adjacent pairwise interaction counts to reflect a merging of adjacent genomic bins. One iteration of this process cuts the dimension of the contact matrix roughly in half. For each matrix generated in the series, we ignore the diagonal elements as we would in the original full contact matrix. Once we generate the multiscale series of matrices, we execute the series of optimizations in the reverse order, beginning with the smallest matrix and ending with the full matrix. We initialize the smallest optimization randomly from a standard multivariate normal distribution, obtain a solution, and then upsample this solution (i.e. interpolate between the solution nodes) to initialize the next larger optimization problem in the series. We continue this iterative process of solving successively larger optimizations, using an upsampled version of the current solution as an initialization to the next higher resolution problem, until we finish with the full solution. SIMBA3D implements this multiscale technique in Python, whereby at each scale the optimization is solved using the BFGS method [7, 16] with analytical gradient (see Supplemental Text for the gradient expression).

Although we solve several optimization problems in the above multiscale approach compared to just one in a standard approach, the computation time is significantly reduced. Since the smaller optimizations are relatively fast compared to the full resolution version, the multiscale approach is essentially a systematic way of cheaply providing a good initialization to the full problem. The combined cost of producing this initialization and executing the full resolution optimization is less than the cost of executing the full resolution optimization with a full resolution random initialization. Moreover, our experimental results show that we achieve on average a lower energy, i.e. better quality, solution using the multiscale approach compared to that of the standard approach of random full resolution initialization. An additional consequence of using the multiscale approach is that by design, the space of obtainable full resolution local solutions is limited by the initial smallest resolution. In extreme cases the very small resolution problems may only have one solution; therefore, if one wishes to explore the local solution space of the full resolution by using different random initializations while still enjoying the benefits of reduced computation time, one must strike a reasonable compromise in initial scale size.

## 3 Results

Here we present some representative results from SIMBA3D applied to the reconstruction of chromosome structures using the mESC dataset [22], with a more extensive presentation of results left for the supplementary material. A complete estimation solution for a simulated configuration, intended as an illustration of the estimation process, is presented in Figure S1 of the supplementary material. To highlight the influence of parameter selection on the results, Figure 1 illustrates the effect of the relative values of the three weights — *λ_1_*, *λ_2_*, and *λ_3_* — on the resulting estimated structures. As expected, higher values of these weights lead to increases in the respective properties they emphasize. For instance, an increase in *λ_3_* leads to the chromosome structure bearing more resemblance to the structure estimated from the bulk data alone. Figure S2 in the supplementary material provides a much more extensive listing of such results for several single cells under a broad range of parameter values.

**Figure 1:**
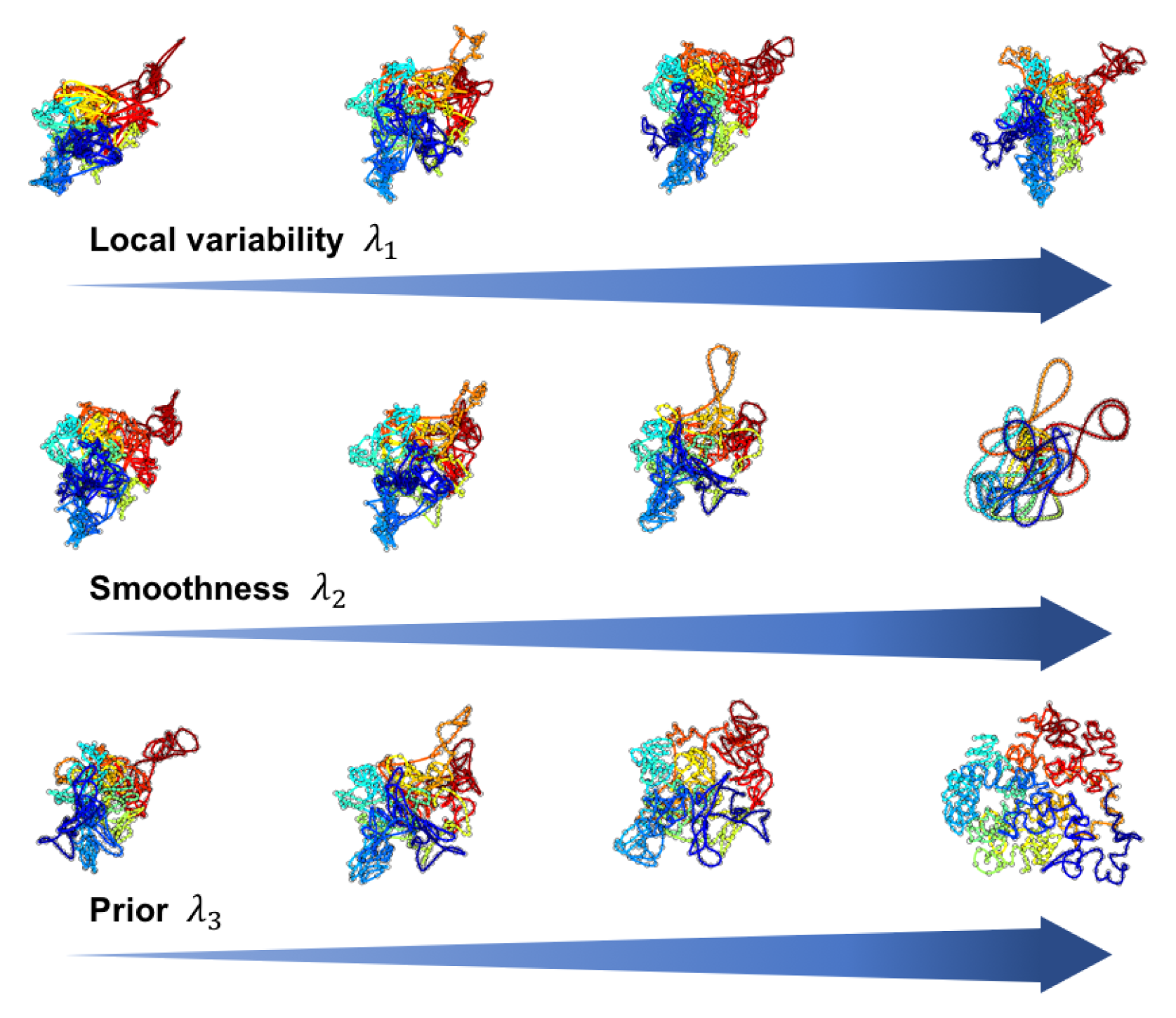
Effect of model parameters on 3D reconstruction quality. We compute the 3D reconstruction as a function of parameter values using the Hi-C data matrix associated with chromosome 19 in cell 1 of the mESC dataset from [22]. The top row of structures from left to right shows the effect of an increased weight *λ_1_* on the parameterization penalty *h*_1_. We vary *λ*_1_ = 0.01, 0.1, 1, 10 and fix *λ*_*2*_ = *λ*_3_ = 0 to obtain four solution curves with exponentially increasing penalty weight. The center row of structures from left to right shows the effect of an increasing weight *λ_2_* on the smoothing penalty *h*_2_. Here, we vary *λ*_2_ = 0.01, 0.1, 1, 10 and fix *λ_1_* = 0.5 and *λ_3_* = 0 to obtain these four solution curves. Finally to show the effect of incorporating the bulk data – the mESC chromosome 19 population matrix – in the analysis, we vary *λ_3_* = 0.01, 0.1, 1, 10 and fix *λ*_1_ = 0.5, *λ*_2_ = 1 to obtain the four structures on the bottom row. We computed all structures using the multiscale approach with *n* = 73, 146, 292, 584.

Figure 2 studies the nature of solutions resulting from different initializations on the same data. Due to the vast search space in which the structure estimation is performed as well as the non-convexity of the objective function, the optimization procedure in SIMBA3D cannot ensure convergence to a *unique* global solution. Instead, the output structure represents one of many different local optima that can be reached depending on the initialization. Despite the existence of several local optima, multiple configurations resulting from the same cells do in fact cluster together in the shape space, as illustrated using a pairwise RMS distance matrix and dendrogram in this figure. The clustering observed here lends further validity to the inferred structures.

**Figure 2:**
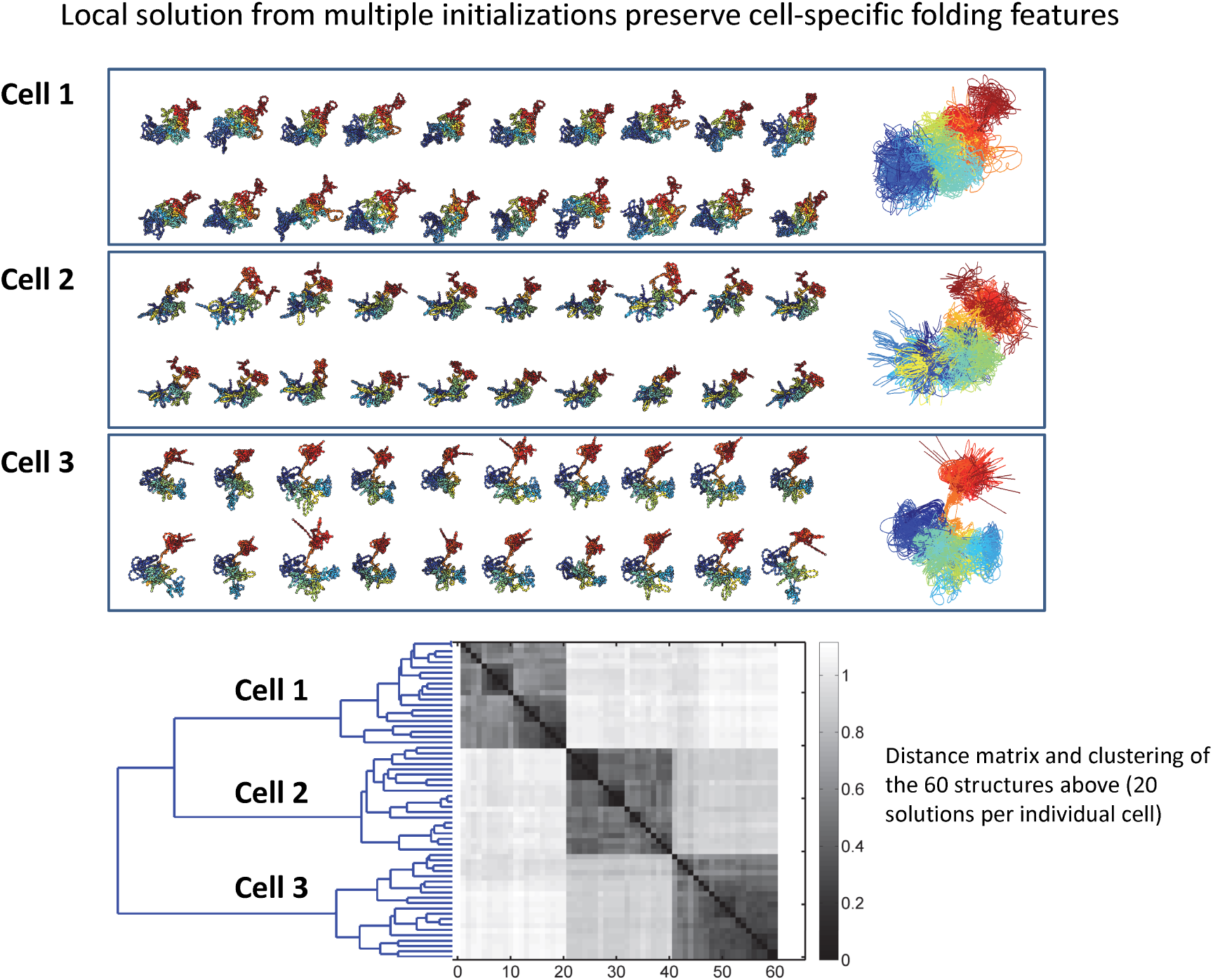
Similarity between ensembles of solutions across cells. Here we show twenty local solutions obtained for chromosome 19 in each of three cells – cell 1, cell 2, and cell 5 – in the mESC dataset using fixed *λ* = (0.5, 1, 0.1). We computed all structures using the multiscale approach with *n* = 73, 146, 292, 584, and for each cell we used the same twenty random initializations at the smallest scale. All displayed solutions are rotationally aligned. For each cell we show the twenty obtained solutions separately, and additionally, to help visualize the variability inherent to the local solutions within cells, we plot the three groups of solutions on top of each other in three respective windows. We then show the clustering of all 60 solutions in shape space via a 60 *×* 60 pairwise RMS distance matrix andassociated dendrogram plot.

The use of the multiscale optimization technique is beneficial for several reasons. It first estimates broader, coarser structures and then adds smaller details, thereby avoiding the abundance of local traps present at the highest resolution. In addition to reaching a lower energy solution on average, it also speeds the algorithm significantly due to low-dimensional searches in the early stages. Figure 3 quantifies gains in computational cost and final energies due to this multiscale approach. Finally, Figure S3 in the supplemental material displays estimated configurations for all 20 chromosomes in all 8 cells present in this mouse dataset using the multiscale approach and fixed penalty weights.

**Figure 3:**
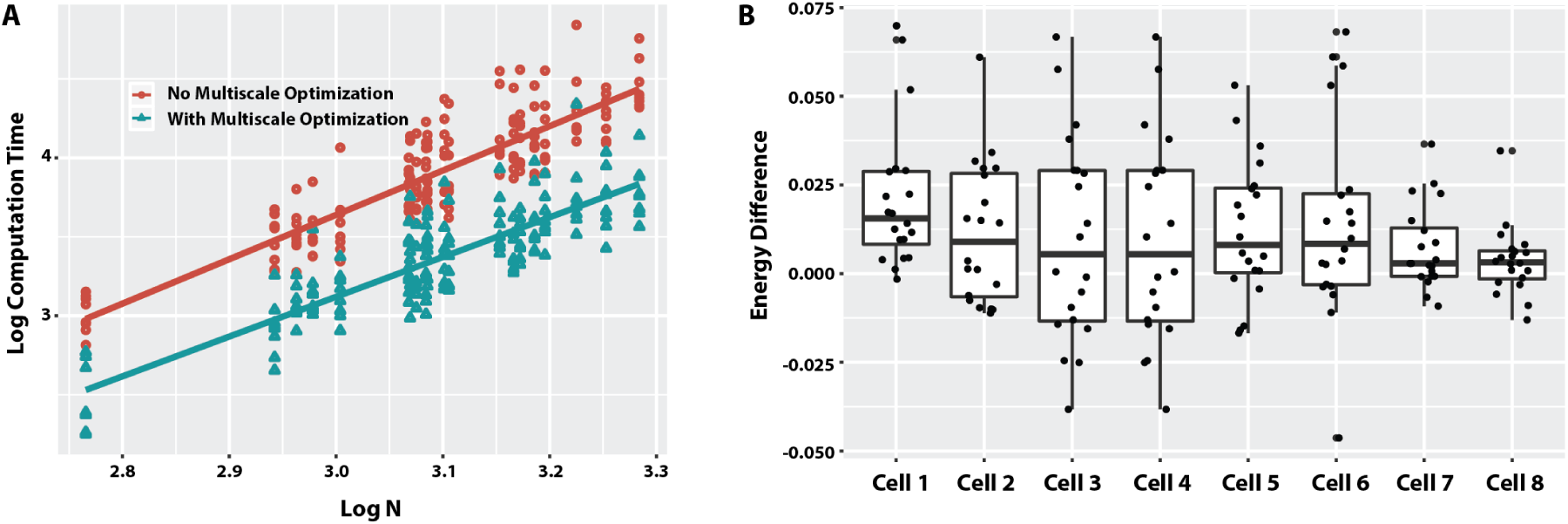
Improvement in computation time and final energy with multiscale optimization. Comparison of results with and without the multiscale approach executed on all 20 chromosomes in each of the 8 cells available in the mESC dataset. (a) The log scale computation time is plotted over log scale number of nodes with regression lines fitted. (b) Boxplot of the difference in final energies obtained for each of the 8 cells. The difference is computed by subtracting the energy obtained via the multiscale approach from the energy obtained without the multiscale approach. That is, a positive number here indicates that the multiscale approach yielded a lower energy solution.

An illustration of this method can be seen in the supplemental movie MovieS1.

## 4 Discussion

SIMBA3D is a Bayesian framework for estimating 3D chromosome structures from single-cell Hi-C data, using penalties for regularization of the estimated structures and using additional information from the bulk Hi-C data. Using multiscale optimization tools and a BFGS routine, it generates computationally efficient inferences and compares these across different initializations and different data (cells). Clustering of solutions in the shape space from the same cell data supports the validity of these solutions.

## URL

The SIMBA3D software is available at https://github.com/nerettilab/SIMBA3D

## Acknowledgments

This work was supported by the NIH Common Fund Program, grant U01CA200147, as a Transformative Collaborative Project Award (TCPA) to TCPA-2017-NERETTI to NN and AS. Additionally, we acknowledge the support of Dr. Frank Crosby at the Naval Surface Warfare Center Panama City Division for funding MR and DB through the In-House Laboratory Independent Research (ILIR) program.

## 5 Authors’ contributions

Conceived the study: NN and AS. Developed the methodology: NN, AS, FH, DB, and MR. Performed the computational analysis: DB, MR, SE. Contributed to the supervision: NN, AS, and FH. Wrote the manuscript: NN, AS, FH, DB, and MR.

## SUPPLEMENTAL TEXT

### 1 Details of Methodology

Suppose that the genome is partitioned into *n* equally sized, disjoint segments, or bins. Let *C* be the *n* × *n* data matrix obtained from a Hi-C experiment. The *ij*’th entry of *C*, call it *c_ij_*, represents the number of observed interactions between segments *i* and *j*, and thus, *C* is naturally a symmetric matrix. Let *x_i_* ∈ ℝ^3^ be the center of mass of the *i*th segment, and let *X* ∈ ℝ^*n*×*p*^ be the collection of all such *x_i_*’s. The problem at hand is twofold: (1) to estimate the structure *X* from the Hi-C data matrix, and (2) to compare the shapes of multiple structures obtained either from the same or different data matrices. In both the estimation and analysis stages, we consider a structure *X* to be equivalent modulo scale, translation, and rigid rotation/reflection.

#### 1.1 Structure Estimation

Varoquaux *et al* [28] link the *ij*’th interaction count with the *ij*’th pairwise distance via the following probability model:

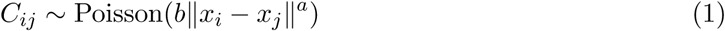
for *j > i*, with ‖ · ‖ being the standard Euclidean norm, and for scalars *a <* 0 and *b >* 0. Since *a <* 0, the expected number of interactions between segments *i* and *j* is larger when the segments are located closer together in space, and this expected number behaves according to a power law with power *a*. Varoquaux *et al* [28] derive the theoretically optimal value of *a* = *−*3 from principles of polymer physics. Furthermore, the parameter *b* acts as a scaling parameter.

The probability mass function for the Poisson random variable given in Eq. 1 is given by

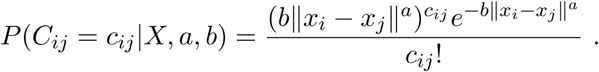

Thus, given a Hi-C matrix with independent entries, the log-likelihood function is written as

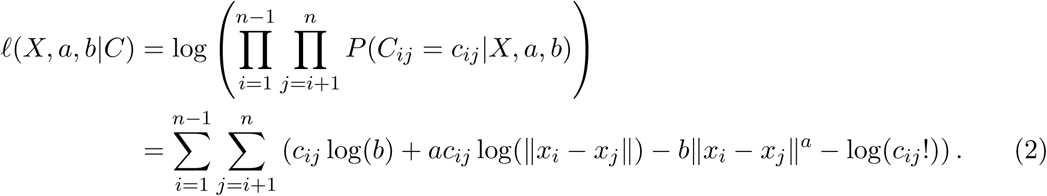

From the log-likelihood function above, one can see that for a given value of *a*, the parameter *b* is non-identifiable because *ℓ*(*X, a, γ^a^b|C*) = *ℓ*(*γX, a, b|C*) for any scalar *γ >* 0. That is, changing *b* is equivalent to changing the scale of *X*, and since *X* is considered equivalent modulo scale, the choice of *b* is arbitrary. Define the function *g* as the negative log-likelihood function, dropping terms that are constant with respect to *X*, i.e.,

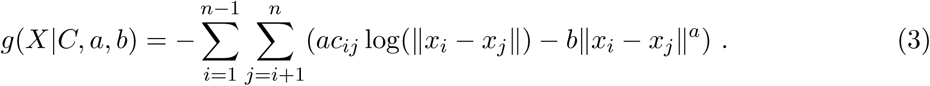

In structure estimation, we consider the parameters *a* and *b* to be fixed and known values; thus, we include them with the data *C* as given when writing the function *g*. The maximum likelihood estimate (MLE) of *X* is computed as the minimizer of *g*; that is,

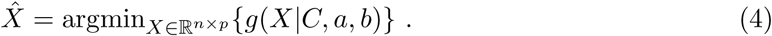

In situations when the data matrix *C* is sparse or noisy, the standard MLE in Eq. 4 can be biologically unrealistic or even fail to converge. Thus, we design several additive penalty terms to regulate the maximum likelihood solution, which leads us to the following penalized likelihood-based objective function:

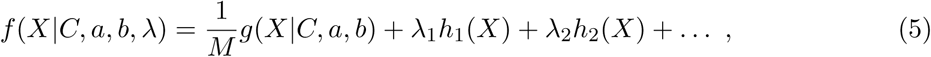
where *M* =∑ _*j>i*_ *c_ij_* is the sum of the upper triangular entries of *C*, *h_i_*(*X*) is the *i*th penalty function, and *λ* = (*λ_1_*, *λ_2_*, *…*) is the vector of penalty weights with *λ*_*i*_ ≥ 0. Wenormalize the function *g* by *M* so that the effects of the data and the penalty terms are in the same proportion for different datasets. We estimate the structure *X* by performing an unconstrained minimization of Eq. 5; that is,

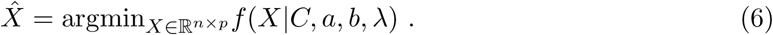

The choice of the penalty weights *λ* is left to the user and can influence the solution greatly.

#### 1.2 Penalty Terms

##### 1. First Penalty

Define the first penalty as

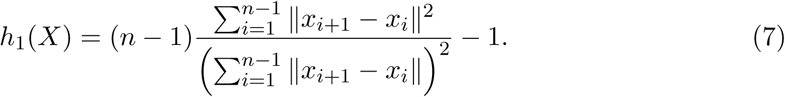

The interpretation of *h*_1_ is the following. Define 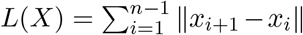 as the length of *X*, and let 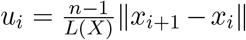 be the distance between the *i*th pair of adjacent points in *X*, when *X* has been rescaled to have length *n* − 1. Then, one can show that 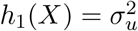, where 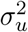 is the variance of the *u_i_*’s, and therefore, the effect of the penalty *h*_1_ is to reduce the variability of the distances between adjacent points of *X*. The configuration which minimizes *h*_1_(*X*) is such that all the *u_i_*’s are equal to 1, i.e. adjacent points in *X* are all the same distance apart. The minimum value of *h*_1_ is 0 regardless of the value of *n*. Furthermore, notice that since *h*_1_(*γX*) = *h*_1_(*X*) for any *γ >* 0, *h*_1_(*X* + *y*) = *h*_1_(*X*) for any *y* ∈ ℝ^*p*^, and *h*_1_(*XR*) = *h*_1_(*X*) for any *p* × *p* orthogonal matrix *R*, the penalty *h*_1_ is invariant to scale, translation, and rotation/reflection.

##### 2. Second Penalty

Define the second penalty as

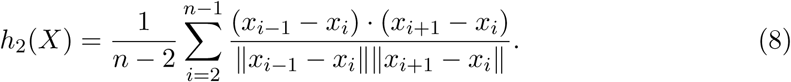

The interpretation of *h*_2_ is the following. If *θ_i_* is the angle created by the triplet of points (*x_i_*_−1_*, x_i_, x_i_*_+1_), and *y_i_* = cos(*θ_i_*), then *h*_2_(*X*) =*y̅*, the sample mean of all the *y_i_* ‘s. Therefore, the minimizer of *h*_2_ is such that cos(*θ_i_*) = −1 for all *i* = 2 …, *n* − 1. This occurs when *X* is a straight line with all *θ_i_* = *π*; hence, the effect of the penalty *h*_2_ is to enforce a level of smoothness to *X*. The penalty *h*_2_ has a minimum value of −1 regardless of the value of n and is invariant to scale, translation, and rotation/reflection.

##### 3. Bulk Prior

Suppose further that another data matrix *C′* is available to us and represents the collective results of prior Hi-C experiments. For example, in the case of a single cell Hi-C matrix *C*, we could also have available to us bulk Hi-C data from the same type of cell, or, alternatively, *C^′^* could be equal to the sum of several other single cell Hi-C matrices. Define the bulk penalty *h*_3_ as the function *g* evaluated using *C′* and normalized by 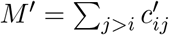. That is, let

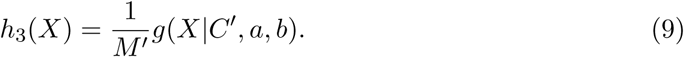

Notice that if we let 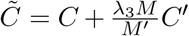 and 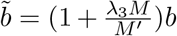, then the term *g*(*X|C, a, b*)*/M* + *λ_3_h*_3_(*X*) is equal to *g*(*X C̃, a,b̃*)*/M* Therefore, the effect of *h*_3_ is essentially to add a scalar multiple of *C′* to the data *C* and perturb the parameter *b*. If *λ_3_* is chosen to be small enough, then this penalty term will only slightly alter *C* and not overwhelm the original data with the bulk data. If *C* is sparse and *C′* is not, then the addition of this penalty term with a small enough *λ_3_* eliminates the sparseness of *C* by replacing many of the 0 entries with small numbers that are biologically more meaningful than random noise.

### 2 Details of Optimization Procedure

There are many possibilities for penalty terms, and thus we write the objective function in Eq. 5 in generality to include any number of penalty terms. However, for practical implementation of the optimization problem in Eq. 6, we use only the penalties *h*_1_, *h*_2_, and *h*_3_ defined in Eqs. 7, 8, and 9 for three reasons. First, each penalty term has a straightforward and biologically meaningful interpretation. Second, the formula for each term is relatively simple and inexpensive to compute – in particular, since *h*_3_ becomes absorbed into the function *g*, the addition of this penalty requires an essentially zero increase in overall computation time. Third, along with the function *g*, we can write an analytical expression for the gradient of each penalty term, and therefore, we can write a gradient expression for *f*. By inputting the expression of ∇ *f* to a numerical solver, we can maintain computational tractability on a personal computer for the large values of *n* typically seen in real data sets. Using these three penalty functions, *f* can be written as

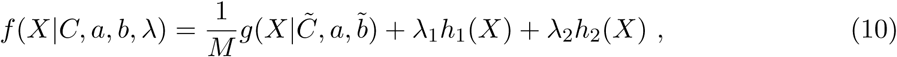
where *C ̃* and *b̃* are defined in the text following Eq. 9, and we consider this objective function for the remainder of this work.

#### 2.1 Gradient Expressions

In order to decrease the computation time when evaluating Eq. 6, it is often helpful to provide the analytical expression for ∇*f* to a numerical optimizer, where the objective function *f* is defined in Eq. 10. The gradient of *f* at *X* is written as

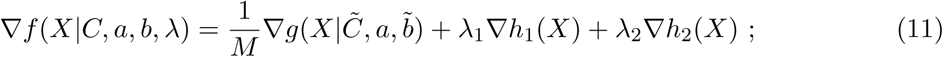
therefore, in order to build the expression for ∇*f*, we need to compute the expressions for ∇*g*, ∇*h*_1_, and ∇*h*_2_, where *g*, *h*_1_, and *h*_2_ are defined in Eqs. 3, 7, and 8, respectively.

For each term in *f*, we compute the expression for the partial derivative at *x_i_* ∈ ℝ^*p*^ for each *i* = 1…, *n*. The *i*th partial derivative evaluated at *X* is a vector in ℝ^*p*^, and thus, the full gradient evaluated at *X* is a member of ℝ^*n×p*^. The *i*th partial derivative of *g* is given by

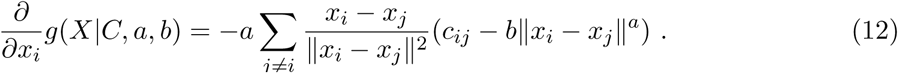

Before computing the partial derivatives of *h*_1_ and *h*_2_ it is useful to define *d_i_* = *x_i_*_+1_ − *x_i_* and *w_i_* = ‖*d_i_*‖ for *i* = 1…*, n −* 1. Also, let 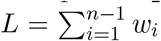 and 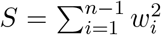 Now, the *i*th partial of *h*_1_ *for i* = 2, …, n − 1 can be written as

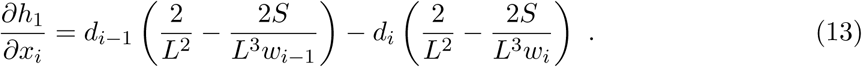

The remaining endpoint partial derivatives for *h*_1_ are given as

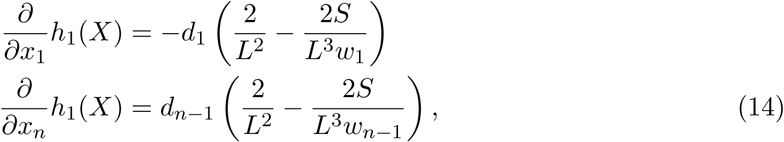

The *i*th partial derivative for *h*_2_ is a bit trickier to compute than that of *g* and *h*_1_ since it requires multiple applications of the chain rule, product rule, and quotient rule; nevertheless, the resulting expressions are still manageable. For *i* = 3,…, *n* − 2, we compute the *i*th partial as

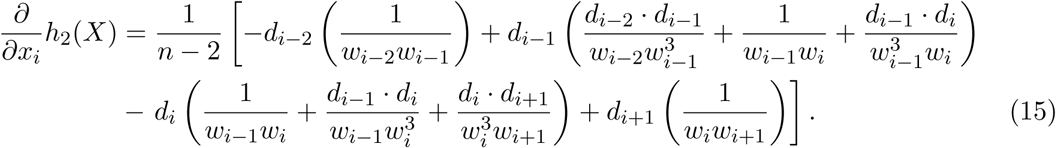

The remaining four endpoint partial derivatives are as follows:

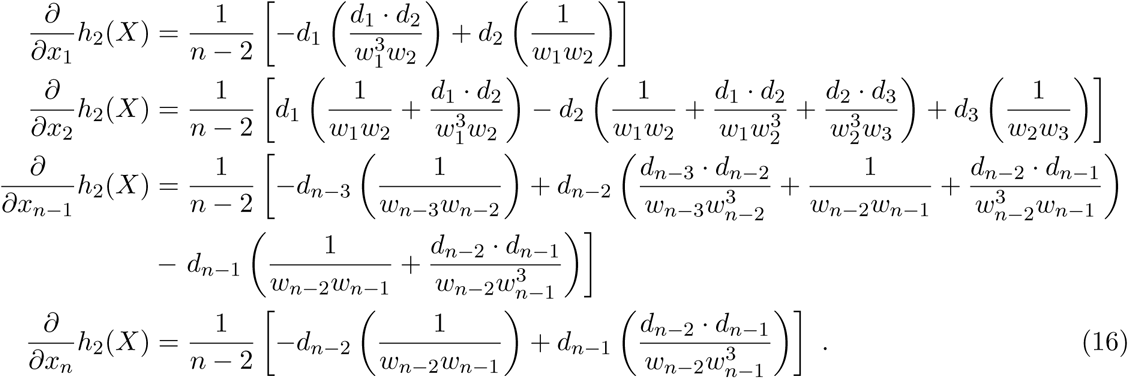
Now, using Eqs. 12, 13, 14, 15, and 16, we can write the *i*th partial of *f* as

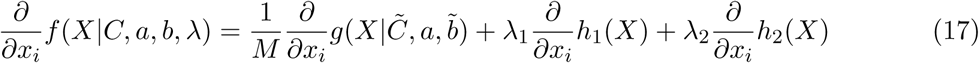
and thus, using Eq. 17, we write the gradient of f in Eq. 11:

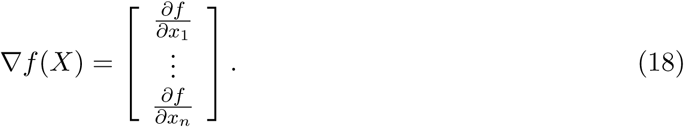

#### 2.2 Software Implementation and Optimization Techniques

Although most scientific computing platforms provide several built-in unconstrained optimization algorithms to solve such problems, we have selected Python as the programming language for SIMBA3D due to its ease of use across a broad scientific community. It is becoming widely adopted in the field because it is open source and does not require a proprietary license to run. Also, specifically addressing the problem at hand, Python ‘s “scipy” package has a wealth of mature built-in scientific computing functions that are necessary for solving this type of optimization problem. In particular, the “scipy.optimize” module offers several popular options for unconstrained optimization algorithms, including Nelder-Mead, Powell‘s method, BFGS, Conjugate Gradient, etc. Python thus gives us the most flexibility and usability for the SIMBA3D code package.

After testing Python‘s built-in algorithms on various simulated and real datasets, we found the quasi-Newton methods BFGS and L-BFGS using our analytical gradient to yield the best performance in terms of numerical stability, computational efficiency, and solution quality when optimizing Eq. 6. With BFGS, the inverse Hessian operation is approximated recursively for each iteration, and for high dimensional optimizations (like for 3D chromosomal reconstruction) this procedure can become memory intensive. With the limited memory version L-BFGS, the approximated inverse Hessian operation depends on values stored from a fixed number of previous iterations. For details see [11, 3, 16, 7]. In our experience with applying these quasi-Newton methods specifically to genome architecture reconstruction, we found that BFGS more reliably finds solutions with slightly smaller energies, i.e. better quality solutions in terms of our objective function, than the L-BFGS. However, the reduction in computation time and memory offered by the the limited memory version makes it a competitive alternative.

In addition to Python ‘s built-in optimization routines, we also tested our own implementation of Nesterov ‘s accelerated gradient method [15]. This method is a simple and elegant modification of the standard gradient descent that uses an additional “momentum “ term to achieve the theoretically optimal convergence rate for a first order method. It has enjoyed a recent surge in popularity due to the publication of a series of physics-based convergence proofs [29, 23, 30] that are more widely relatable and understandable than Nesterov‘s original proofs. In many test cases we saw a significant computational speedup over the BFGS, and we see great promise in this optimization method in the future. Unfortunately, since the method is not a relaxation scheme, i.e. it does not necessarily decrease the objective function value after each iteration, we found it difficult to devise an appropriate stopping criteria that would generalize to all data sets. Another difficulty that arose in our implementation of this algorithm was an automatic step size selection routine via backtracking, and thus achieving automated numerical stability for a general data set proved to be a challenge. We believe we can overcome these challenges to implementing this method, but in the interest of time and to remain within the scope of this research, we feel that BFGS performs well enough to use for SIMBA3D, especially when combined with the multiscale approach detailed in the main text.

### 3 Additional Results

Here we present a series of results that are not shown in the main text, which come from three computational experiments. The first experiment is carried out on simulated data and shows how the computation time increases as the number of nodes *n* increases. In the second experiment we execute an exhaustive parameter search by running the optimization with various settings of the penalty weights *λ* on Chromosome 19 of each of the eight single cells in the mESC data set. Finally, in the third experiment we compare the performance of the BFGS and L-BFGS with and without the multiscale approach in terms of computation times and final energies on all 20 chromosomes and all 8 cells in the data set. Fig. 3 in the main text summarizes the results of this experiment, but here we present the results in their entirety.

Table S1 and Fig. S1 summarize the results from the simulated computation time experiment. The ground truth curve is a unit length double spiral shape as seen in Fig. S1 (A). For each row of Table S1, the number of nodes in the double spiral curve was doubled using upsampling via spline interpolation, and a corresponding data matrix was generated from the upsampled curve. Fig. S1 (B) shows an example data matrix that corresponds with the curve and number of nodes presented in Fig. S1 (A). The entries of the upper triangular portion of each data matrix were simulated using the independent Poisson random variables *C_ij_* ~ *Poisson*(*b*‖*x_i_* − *x_j_*‖*^a^*) with *a* = −3 and *b* set large enough to avoid simulating a sparse matrix. For each number of nodes *n*, ten initializations to the optimization procedure were generated as random samples of size *n* from the 3-dimensional standard multivariate normal distribution. Then for each number of nodes and for each initialization, the optimization was run using the BFGS without the multiscale approach, and the computation time and final energy were recorded in each case. However, in the case with 2560 nodes, the experiment was stopped prematurely after the sixth initialization due to an unreasonably long computation time. The penalty weight vector *λ* remained fixed for all optimizations in the experiment, with *λ_3_* = 0 due to the absence of simulated prior data. Fig. S1 (C) shows the evolution of the energy, i.e. the objective function evaluated at each iteration of the optimization algorithm, for one initialization and using the data given in panel (B), and panel (D) shows the final estimated solution curve. Panel (E) is a scatter plot of the log computation time using each of the ten initializations versus the log number of nodes, including a fitted regression line.

Fig. S2 shows the results of an exhaustive grid search experiment over the penalty weight vector *λ* for Chromosome 19 in each of the eight cells. The purpose of the experiment is to explore how certain combinations of values for *λ* affect the solution properties, which offers the user some guidance on selecting a reasonable value of *λ*. Here, every possible combination of *λ*_*i*_ = 0, 0.001, 0.01, 0.1, 1 for *i* = 1, 2, 3 was set, totaling 5^3^ = 125 different settings. Ten random initializations were generated from the standard multivariate normal distribution for use in each of the 125 cases; thus, 1250 solution curves were obtained for each cell for a total of 8 *×* 1250 = 10000 solutions. The experiment was performed using the BFGS method with and without the multiscale approach.

The results presented in Fig. 3 of the main text have been extended in Fig. S3 and Table S2 to include comparisons using BFGS and the limited memory version L-BFGS with and without the multiscale approach. The experiment shows how computation time varies with respect to the number of nodes by executing the optimization on all 20 chromosomes in each of the eight single cells. It also shows that one can use SIMBA3D on a standard laptop to achieve a full genome reconstruction within a practical time frame. Fig. S3 consists of four appropriately named pdf files (subfigures) that show the final solution curves and energy evolution for each chromosome in each single cell using either BFGS or L-BFGS with and without the multiscale initialization. The penalty weight vector *λ* was fixed throughout the experiment and set to have reasonable non-zero component values according to the results gathered from the previous experiment. Table S2 lists and plots the final computation times and energies from these reports along with other information describing the data.

---

**Figure S1. Example of 3D shape reconstruction in simulated data.** Computation time experiment on simulated data. (A) Ground truth curve using 80 points. (B) Simulated matrix using the Poisson model from Eq. 1. (C) Example of the energy evolution from a randomly initialized curve. (D) Example of a reconstructed curve from the data matrix. (E) Log scale computation time versus the log scale number of nodes plotted for each of the ten initializations.

**Figure S2 (a,b). Model parameter sweep.** This figure consists of two large pdf files that present an exhaustive parameter search on chromosome 19 using the BFGS method with and without the multiscale approach. Figure S2 (a) presents results that were obtained using the multiscale approach, and Figure S2 (b) initializes at the full scale resolution. Each of the three parameters *λ*_*i*_, *i* = 1, 2, 3 varies from 0, 0.001, 0.01, 0.1, and 1.0 to create 5 tables with 5× 5 rows and columns that list the 5^3^ = 125 possible value combinations of *λ* that were tested. For each parameter setting and for each of the eight cells, the algorithm was run with the same set of ten random initializations; thus, this experiment contains a total of 5×5×5×10×8 = 10000 runs for each of the two subfigures. In each panel the ten estimated curves for each setting are rotationally aligned and plotted together, and the iterative energy evolution is plotted below the curves. For the multiscale approach the sequential number of nodes were set as *n* = 74, 146, 292, 584, and the initialized curve of 74 nodes was obtained by downsampling the previously used random full scale initializations. In the energy evolution plots the increasing line thickness indicates the run using *n* = 74, 146, 292, 584 nodes, respectively.

**Figure S3 (a-d). Reconstruction of all chromosomes in each single cell data matrix.** This figure consists of four large pdf files – one for each algorithmic combination of multiscale or full scale, with BFGS or L-BFGS – each showing the result of the optimization executed on all 20 chromosomes for each of the 8 cells with *λ* = (0.5, 1, 0.1). For each chromosome and cell, a single run is collected using a randomly initialized curve. The total computation times and the final energies are displayed below the corresponding curve and energy plot. In displaying the multiscale results, the energy plot contains four evolution curves corresponding to the different multiscale optimizations. For each chromosome the full scale node resolution was roughly halved a total of four times so that the solution was initialized using roughly 1*/*2^4^ the number of nodes in the full scale data, which is different for each chromosome. The thickest and bluest line in the energy evolution plots shows the full resolution run, and the thinner blacker lines show the optimizations using the smaller data matrices.

**Table S1. Computation time summary statistics on simulated data**The mean and variance of the computation times for each number of nodes are computed after executing the optimization using ten different initializations. An example of one run for this experiment is shown in Fig. S1 for the case when there are 80 nodes.

**Table S2. A Comparison of computation times and energies on the full chromosome data.** There are 8 sheets – one for each of the 8 cells – and on each sheet there are 20 rows of data corresponding to the 20 chromosomes. The first 5 columns specify generic information about the single cell data matrix. The following columns correspond to computation times and final energies for single initialized runs using four different methods. A BFGS and L-BFGS label respectively indicates the use of the quasi-Newton BFGS or its limited memory version. The FS tag on the label indicates that the algorithm was initialized on the full scale space. For example with Cell 1 Chromosome 1, the BFGS FS column indicates that BFGS was used and the solution was initialized with a random sample of size 1923 from the standard multivariate normal distribution in ℝ^3^. The MS tag on the label indicates that the curve was estimated using the multiscale approach detailed in the main text. In the multiscale approach the scale was roughly halved a total of four times; thus, for each chromosome the number of nodes in the initialization for the MS case is roughly 1*/*2^4^ that of the FS case.

**Movie S1. Time-course optimization for chromosome 19.** This movie displays the intermediate structures obtained during the optimization procedure. Rotating chromosome structure (tope left). Stationary view of chromosome structure (top right). Energy as a function of iteration step (bottom left). Single cell contact matrix at the current resolution (bottom center). Bulk contact matrix at the current resolution (bottom right).

